# Torsins Organize CLCC1 Assembly to Safeguard ER Bilayer and Lipid Homeostasis

**DOI:** 10.64898/2025.12.14.694184

**Authors:** Yonglun Wang, Renqian Wang, Yuanhang Yao, Rongxian Hou, Lu Yan, Yiechang Lin, Yating Hu, Youlei Li, Lingzhi Wu, Yuangang Zhu, Ning Gao, Chen Song, Xiao Wang, Zhejian Ji, Xiao-Wei Chen

## Abstract

The TMEM41B scramblase and its regulatory partner CLCC1 initiate lipid flux by equilibrating newly-synthesized phospholipids across the endoplasmic reticulum (ER) bilayer, a fundamental process required for diverse events ranging from membrane biogenesis to bulk lipid supply. Loss of CLCC1/TMEM41B causes ER bilayer imbalance, inducing giant ER-enclosed lipid droplets (geLDs) and driving rapid progression into severe MASH. Here we identify CLCC1 as the long-missing client of the lumenal Torsin ATPases, which selectively engage oligomerized CLCC1 at sites of ER bilayer imbalance. Hepatic TorsinA inactivation triggers geLD formation amid disrupted lipoprotein biogenesis and severe MASH, closely phenocopying CLCC1/TMEM41B deficiency. Mechanistically, Torsins act as foldases that drive CLCC1 oligomerization for its recruitment to imbalanced bilayers. Remarkably, ectopic CLCC1 expression reverses cellular and systemic lipid disorders arising from hepatic TorsinA deficiency. Hence, Torsin ATPases emerge as fundamental regulators that organizes CLCC1 and the downstream TMEM41B scramblase to govern lipid partitioning and membrane homeostasis.

## Introduction

The precise coordination of lipid production, storage, and mobilization is vital for cellular and systemic homeostasis^1,2^. Dyslipidemia, resulted from dysfunctional plasma lipid control, represents the major risk factor for cardio-metabolic diseases, so-called No.1 killer^3,4^. Bulk lipids are transported into the circulation via ApoB-containing lipoproteins, initially assembled in the lumen of the endoplasmic reticulum (ER) in metabolically active hepatocytes in the liver or enterocytes in the gut^5,6^. During this process, lipoproteins must acquire substantial amounts of both neutral lipids and phospholipids across the ER membrane, yet the molecular mechanisms governing these trans-bilayer lipid fluxes remain incompletely defined^7–10^. Following their initial assembly, large quantities of lipoproteins exit the ER through the COPII transport machinery, in a selective manner governed by the secretory cargo receptor SURF4 in hepatocytes and enterocytes^11–13^.

The ER is the central hub for lipid synthesis and secretion^14^. While neutral lipids are synthesized between the leaflets of the ER bilayer, phospholipids are asymmetrically produced at the cytosolic leaflet and are subsequently translocated to the luminal leaflet by phospholipid scramblases such as TMEM41B^15,16^. The ER transmembrane protein CLCC1 partners with the TMEM41B scramblase to promote lipid scrambling, thereby licensing lipoprotein biogenesis and the subsequent bulk lipid transport^17^. Loss of CLCC1 or TMEM41B causes catastrophic disruption of lipid flux, leading to giant ER-enclosed lipid droplets (geLDs), severe ER bilayer distortion, and drastically accelerated development of metabolic-dysfunction–associated steatohepatitis (MASH)^17,18^. Intriguingly, CLCC1 dynamically forms oligomers that recognize these imbalanced ER bilayers and facilitate the subsequent TMEM41B recruitment, highlighting phospholipid scrambling as a tightly regulated and dynamic process that is essential for cellular and systemic lipid homeostasis^17^.

Torsins (TorsinA, TorsinB, Torsin2, Torsin3, and Torsin4) belong to the AAA+ ATPase superfamily, which carries out a diverse range of cellular functions^19–28^. However, Torsins represent the only AAA+ ATPases localized within the ER lumen and play key roles in maintaining the integrity of the nuclear envelope, a specialized continuum of the ER^21,23–25,27^. Heterozygous mutation of TorsinA in humans disrupts motor neuron function and causes early-onset dystonia^29,30^, while homozygous loss of TorsinA results in the rarer, much more severe disease arthrogryposis multiplex congenita type 5 (AMC5)^31^. Despite much attention over several decades, the exact molecular function and potential substrate(s) of Torsins remain unclear^19^. Moreover, though Torsins have long been proposed to serve as “unfoldases” to dissemble client protein complexes, these ATPases lack key residues that constitute the central pore required for canonical unfoldase activities^32,33^. Intriguingly, hepatic inactivation of TorsinA disrupts lipoprotein secretion and induces profound MASH pathology in mice, largely resembling the defects triggered by either CLCC1 or TMEM41B loss, though the underlying mechanism of Torsins in lipid control also remains unclear^26^.

Here we show that Torsins act as foldase-like activators of CLCC1 to safeguard lipid homeostasis and ER membrane integrity. Quantitative proteomics identified that Torsins selectively interact with CLCC1 upon the induction of ER bilayer imbalance. Closely recapitulating the unique phenotypes of CLCC1/TMEM41B deficiency, hepatic TorsinA inactivation induces lipid mis-deposition into geLDs and drives pronounced ER bilayer imbalance. At the cellular level, Torsins are required for CLCC1 oligomerization and its recruitment to sites of bilayer asymmetry. *In vitro*, Torsins directly nucleate CLCC1 oligomerization, revealing an unexpected molecular activity for these luminal ATPases. Importantly, ectopic CLCC1 expression restores both cellular and systemic lipid homeostasis in the setting of hepatic TorsinA deficiency, positioning CLCC1 as a principal downstream effector of Torsins in lipid partitioning and membrane homeostasis.

## Results

### Identification of Torsins as inducible partners of CLCC1

Our recent work^17^ led to the hypothesis that CLCC1 oligomerization is a key event in regulating ER bilayer equilibrium and thereby controlling lipid flux. We therefore aimed to identify regulator(s) that drive CLCC1 assembly, using a two-step biochemical screen (Figure 1A). Binding partners of tandem-tagged CLCC1 were first purified from either wild type or TMEM41B knockout (KO) liver, given that loss of TMEM41B strongly enhances CLCC1 oligomerization^17^. Second, quantitative mass spectrometry was employed to identify potential partners that preferentially associate with oligomerized CLCC1. Of note, silver staining revealed a ∼35kD band that selectively co-purified with tagged CLCC1 from TMEM41B-deficient liver, compared to wild type controls (Figure 1B). Quantitative mass spectrometry subsequently identified TorsinA and its paralog TorsinB as selective binding partners of CLCC1 upon TMEM41B deficiency (Figure 1C&D). This inducible interaction was further confirmed by immune-blotting following co-immunoprecipitation (co-IP) between Torsins and endogenous CLCC1 (Figure 1E) or tagged CLCC1 (Figure S1) in murine liver. Reciprocal isolation using ectopically expressed TorsinA also captured elevated interaction with endogenous CLCC1 upon TMEM41B deficiency (Figure 1F).

**Figure 1.**
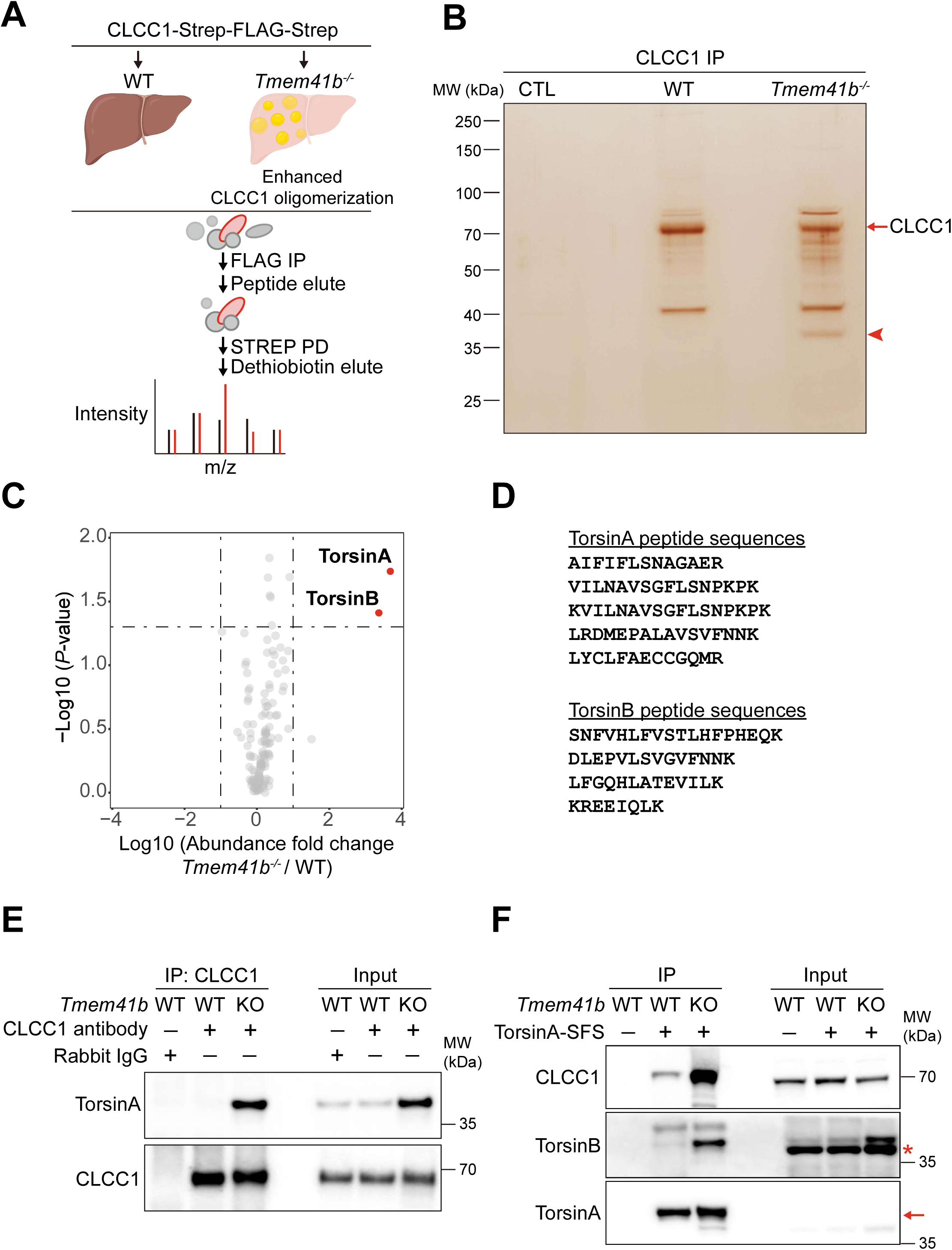
Identification of Torsins as inducible partners of CLCC1. A. Hepatic CLCC1-Strep-FLAG-Strep immune-complexes were first isolated from liver tissues of either wild-type (WT) or liver-specific *Tmem41b* knockout (LKO) mice using anti-FLAG resin, followed by elution with 3×FLAG peptide. The eluted samples were then subjected to the streptactin agarose pulldown and subsequently eluted with dethiobiotin. Finally, the resulting isolates were analyzed by quantitative mass spectrometry (MS). B. Silver staining of the hepatic CLCC1 complex purified from WT or *Tmem41b* LKO mouse livers. Arrows denote the CLCC1 band (top) and putative TMEM41B deficiency-induced partners (bottom), respectively. C. Quantitative proteomics of the hepatic CLCC1 complex isolated from WT (n=3) and *Tmem41b* LKO mice (n=3). D. Peptide sequences of TorsinA and TorsinB detected by MS. E. TMEM41B deficiency induces interactions between endogenous CLCC1 and TorsinA. Immuno-complexes of CLCC1 isolated from murine liver with indicated genotypes were subjected to SDS-PAGE and immunoblotted with the indicated antibodies. The asterisk represents non-specific band. F. Enhanced interaction between ectopic TorsinA and endogenous CLCC1 upon TMEM41B deficiency. Purified products of TorsinA complex from mice livers with indicated genotypes were subjected to SDS-PAGE and immunoblotted with the indicated antibodies. Red asterisk: non-specific band.

### Torsin deficiency induces lipid mis-deposition across the ER membrane

Hepatic inactivation of TorsinA depletes VLDLs and causes drastic hepatic lipid accumulation in mice, although the underlying molecular mechanism remains to be determined^26^. The newly-identified Torsin/CLCC1 interaction prompted us to examine whether disrupted cellular lipid partitioning, a distinct defect upon loss of either CLCC1 or TMEM41B, might underlie the profound pathologies caused by Torsin deficiency. To this end, we inactivated TorsinA, CLCC1, and TMEM41B using CRISPR/Cas9 mediated gene editing in Huh7 cells (Figure S2A-D). Notably, lipid droplets accumulated in TorsinA-deficient Huh7 cells lacked cytosolic LD markers such as Perilipin2, but were instead encircled by the ER membrane marker Cyb5 (Figure 2A&B, upper). The phenotypes closely mimicked lipid mis-deposition into lumenal storages upon CLCC1 inactivation (Figure 2A&B, lower). Electron microscopy revealed that, in contrast to the conventional, cytosolic lipid droplets in wild type hepatocytes, lipid droplets in TorsinA-deficient hepatocytes were enclosed by ER bilayers, confirming their geLD identity (Figure 2C). Moreover, deficiency in either CLCC1 or TMEM41B disrupted cellular localization of NUP358 (Figure S2E), the cytoplasmic filaments of nuclear pore complex (NPC), resembling the NPC biogenesis defect induced by TorsinA inactivation as previously reported^21,24,27^.

**Figure 2.**
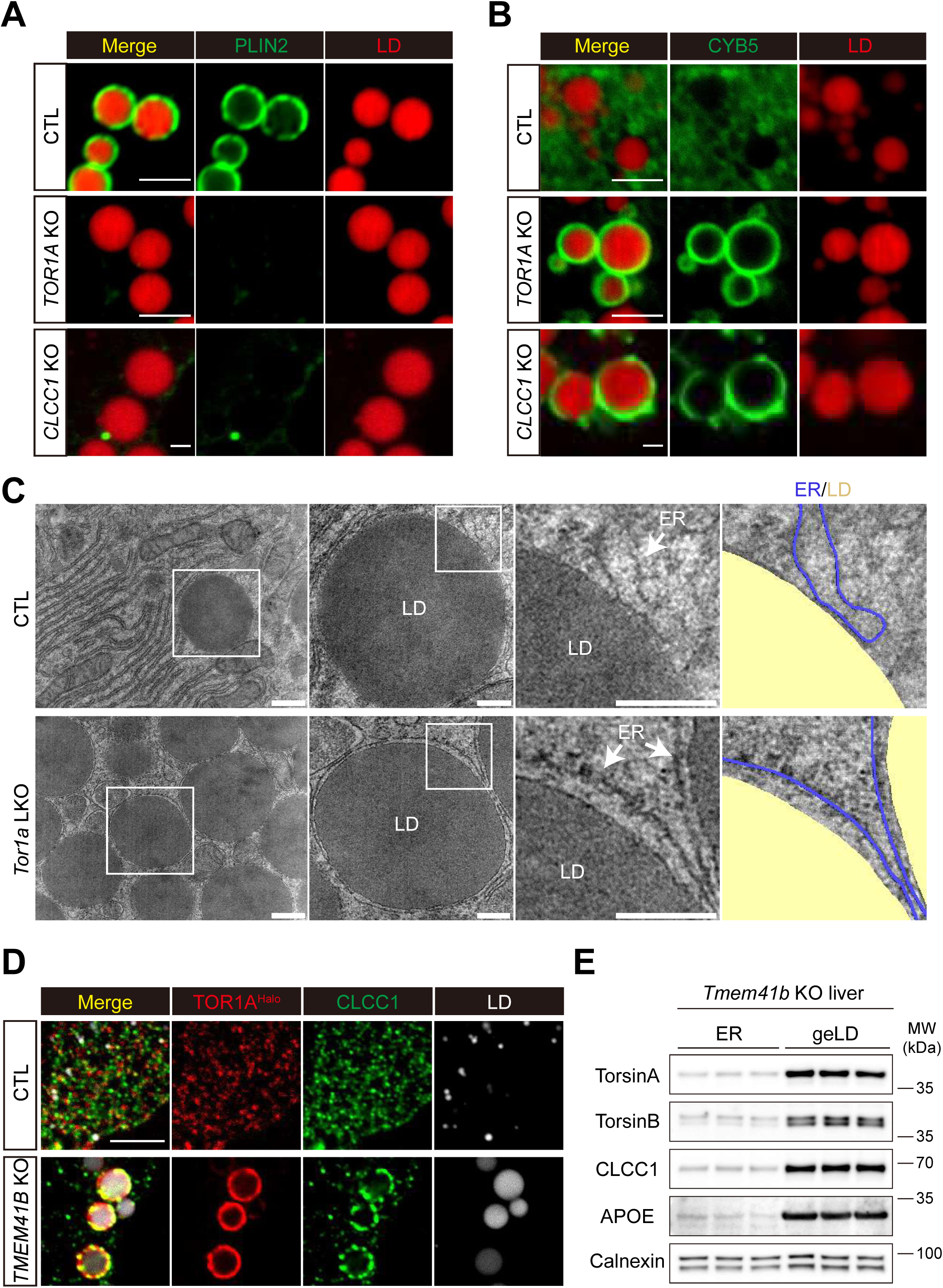
Torsin deficiency induces lipid mis-deposition across the ER membrane. A. The cytosolic lipid droplet (LD) marker PLIN2 is absent from LDs that accumulate in TorsinA-deficient Huh7 cells. Control (top), *TOR1A* KO (middle), and *CLCC1* KO Huh7 cells were infected with AAV-CAG-GFP-PLIN2, fixed, and imaged by confocal microscopy. LipidTOX (red) marks LDs. Representative images of three independent experiments are shown. Scale bar, 2 μm. B. The ER membrane marker CYB5 is enriched in the LDs that accumulate in TorsinA-deficient Huh7 cells. Control (top), *TOR1A* KO (middle), and *CLCC1* KO cells were infected with AAV-CAG-GFP-CYB5 and imaged by confocal microscopy. LDs are stained with LipidTOX (red). Representative images of three independent experiments are shown. Scale bar, 2 μm. C. TEM analysis of the liver ultrastructure in wild-type (WT, top) and *Tor1a* KO (bottom) mice. In the right panels, ER regions are pseudocolored in blue and LDs are highlighted in yellow. Scale bars: 500 nm (left), 200 nm (middle and right). D. TorsinA is enriched in the imbalanced membrane associated with geLDs. Control (top) and *TMEM41B* KO (bottom) Huh7 cells were co-stained with HaloTag JFX554 ligand, anti-CLCC1 antibody, and LipiBlue prior to confocal imaging. Scale bar: 5 μm. E. Enrichment of Torsin proteins in geLDs from *Tmem41b* KO mouse livers. ER and geLD fractions were isolated as per Methods, with equal protein loading across samples. The blot is representative of three independent experiments.

To visualize endogenous TorsinA, we introduced a twin-Strep-Halo tag into the C-terminus of TOR1A by CRISPR/Cas9-mediated homologous recombination in Huh7 hepatoma cells (Figure S3A-C). Consistent with the recruitment of CLCC1 to curved ER membranes enclosing geLDs, TorsinA was similarly enriched at these sites of unmatched membrane bilayers (Figure 2D). Biochemical isolation confirmed the co-enrichment of Torsins and CLCC1 in isolated geLDs (Figure 2E, quantified in Figure S3D), further supporting the role of Torsins in controlling lipid partitioning across the ER bilayer.

### Torsin regulates cellular CLCC1 targeting to imbalanced ER bilayers

The spatial enrichment of endogenous Torsin and CLCC1 at the curved, imbalanced ER bilayer surrounding geLDs prompted us to investigate a potential hierarchy in their actions. The TorsinA^Halo^ Huh7 was employed to track the enrichment of TorsinA and CLCC1 in geLDs. We then acutely inactivated the TMEM41B scramblase by CRISPR/Cas9 to facilitate the monitoring of the early phases of ER bilayer imbalance during geLD induction. Compared to the diffusive TorsinA and CLCC1 distribution in wild type Huh7 cells, loss of TMEM41B triggered their relocations to the ER membranes enclosing the geLDs (Figure 3A). During the early phase of TMEM41B inactivation, a greater percentage of the geLDs were decorated by TorsinA than by CLCC1 (Middle panels, Figure 3A, quantified in 3B). Prolonged TMEM41B inactivation eventually resulted in TorsinA/CLCC1 enclosure of nearly all geLDs (Lower panels, Figure 3A, quantified in 3B). These imaging data therefore indicate that Torsins likely acts upstream of CLCC1. Interestingly AlphaFold3-assisted structural analysis predicted that Torsin oligomers might be encircled by the CLCC1 oligomer and occupy the central cavity of the latter assembly (Figure 3C). Concerning the promoting effects of CLCC1 in TMEM41B-mediated lipid scrambling^17^, we further stimulated the interplays between TorsinA/CLCC1 complex and membrane lipids. We carried out triplicate 5 μs coarse grained molecular dynamics simulations of the complex embedded within a DOPC membrane (Figure 3D). These simulations revealed decreased lipid order (Figure 3E, left) around the CLCC1 oligomer, primarily corresponding to the tilting of the lipid molecules perturbed by the proteins (Figure 3E, right), indicating that oligomerized CLCC1 may locally destabilize the bilayer.

**Figure 3.**
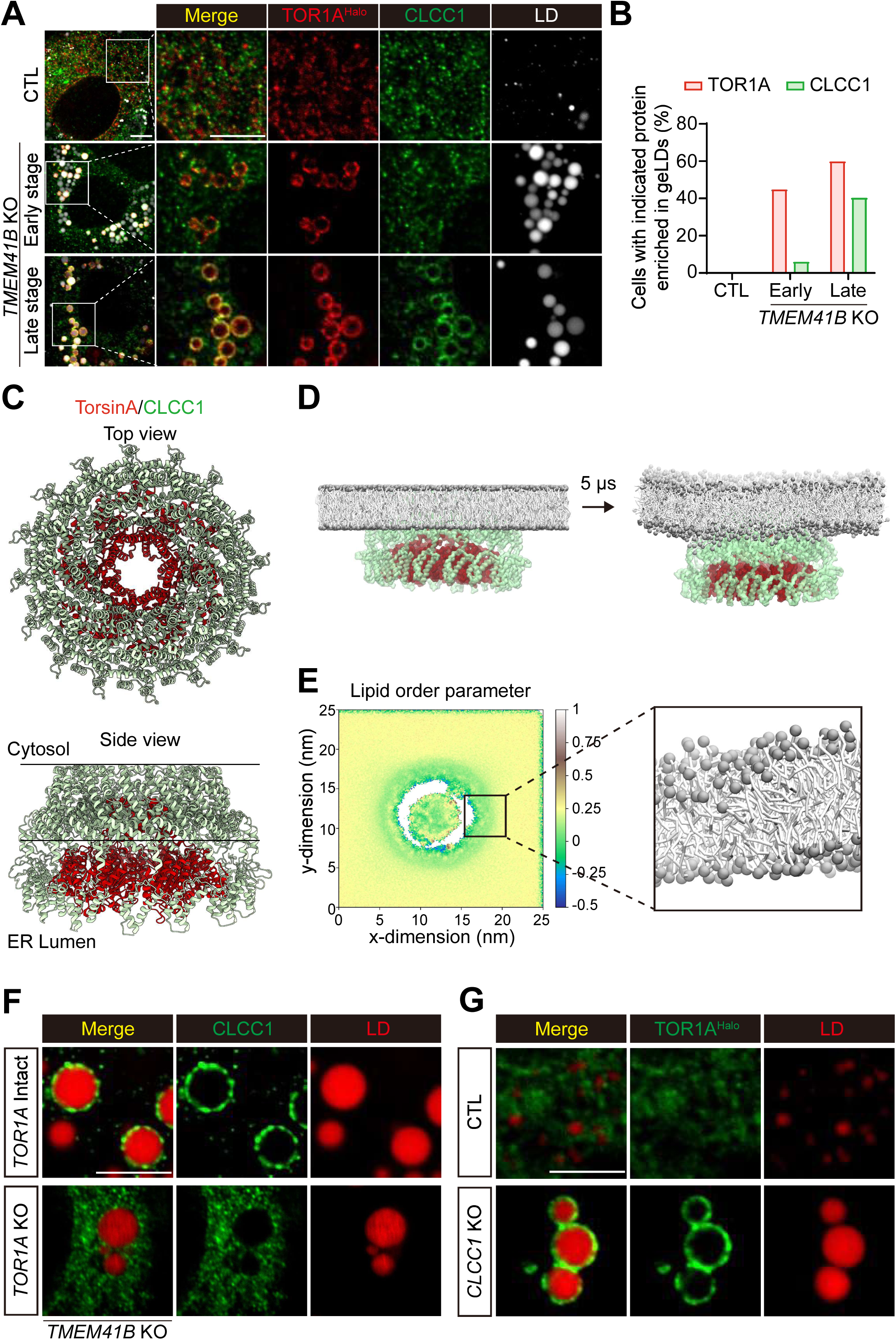
Torsin regulates cellular CLCC1 targeting to imbalanced ER bilayers. A. Hierarchical recruitment of TorsinA-Halo and CLCC1 at geLDs. TOR1A-Halo knock-in Huh7 cells stably expressing spCas9 were infected with lentiviral sgRNAs targeting *LacZ* (control) or *TMEM41B*. Cells were fixed and co-stained with HaloTag JFX554 ligand and the anti-CLCC1 antibody at certain time points (described in Method), followed by confocal microscopy. Lipid droplets were visualized with LipiBlue. Images are representative of three independent experiments. Scale bar: 5 μm. B. Quantification of geLD targeting from (A). Recruitment was quantified as the percentage of cells exhibiting clear enrichment of the indicated proteins (TorsinA-Halo or CLCC1) at geLDs. C. AlphaFold3 prediction of the TorsinA oligomer encircled by the CLCC1 oligomer. D. Coarse-grained MD simulation of the TorsinA/CLCC1 complex in a phospholipid bilayer. Depicted are the initial system setup (left) and the final conformation after 5 µs of simulation (right). E. The TorsinA/CLCC1 complex induces membrane lipids disorder. (Left) Lipid acyl chain order parameters were quantified from the simulations in (D). (Right) Visualization of the simulation snapshot reveals the pronounced tilting of lipids proximal to the protein complex. F. TorsinA is required for CLCC1 recruitment to geLDs. Compared to *TMEM41B* KO Huh7 cells, CLCC1 recruitment to geLDs was abolished in *TMEM41B* & *TOR1A* double KO cells. Cells were co-stained with anti-CLCC1 antibody and LipiBlue, then imaged by confocal microscopy. Images are representative of three independent experiments. Scale bar: 5 μm. G. TorsinA recruitment to geLDs is independent of CLCC1. Confocal microscopy of control and *CLCC1* KO *TOR1A*-Halo knock-in Huh7 cells shows that TorsinA-Halo (labeled with JFX554 ligand) localizes to geLDs in both genotypes. Lipid droplets were stained with LipiBlue. Images are representative of three independent experiments. Scale bar: 5 μm.

We further examined the potential hierarchical targeting of cellular Torsins and CLCC1. Indeed, while CLCC1 was enriched at geLD-surrounding membranes upon TMEM41B inactivation, such recruitment was abolished by additional inactivation of TorsinA (Figure 3F). By contrast, TorsinA could translocate to geLD-surrounding membranes in CLCC1-deficient cells (Figure 3G), as previously seen with TMEM41B-deficient cells (Figure 2D). Taken together, the imaging and biochemical data revealed that Torsins act upstream of CLCC1 to control CLCC1 targeting in cellular settings.

### Torsins act as protein foldases to promote CLCC1 oligomerization in lipid control

The oligomerization-dependent recruitment CLCC1 to the imbalanced ER bilayers^17^ and the Torsin-dependence of the process (Figure 3E) suggest the intriguing possibility that, in contrast to the long-held hypothesis that Torsins are unfoldases to disassemble client substrates, these unique AAA+ ATPase may instead promote high-order *<u>assembly</u>* of specific clients. To test this new hypothesis, we first sought to map the domains within CLCC1 responsible for Torsin interaction (Figure 4A). Consistent with the lumenal orientation of Torsins, the lumenal domain of CLCC1 (residues 19-184) was required for Torsin interaction (Figure 4B). Notably, this region was also required for the cellular oligomerization and function of CLCC1^17^.

**Figure 4.**
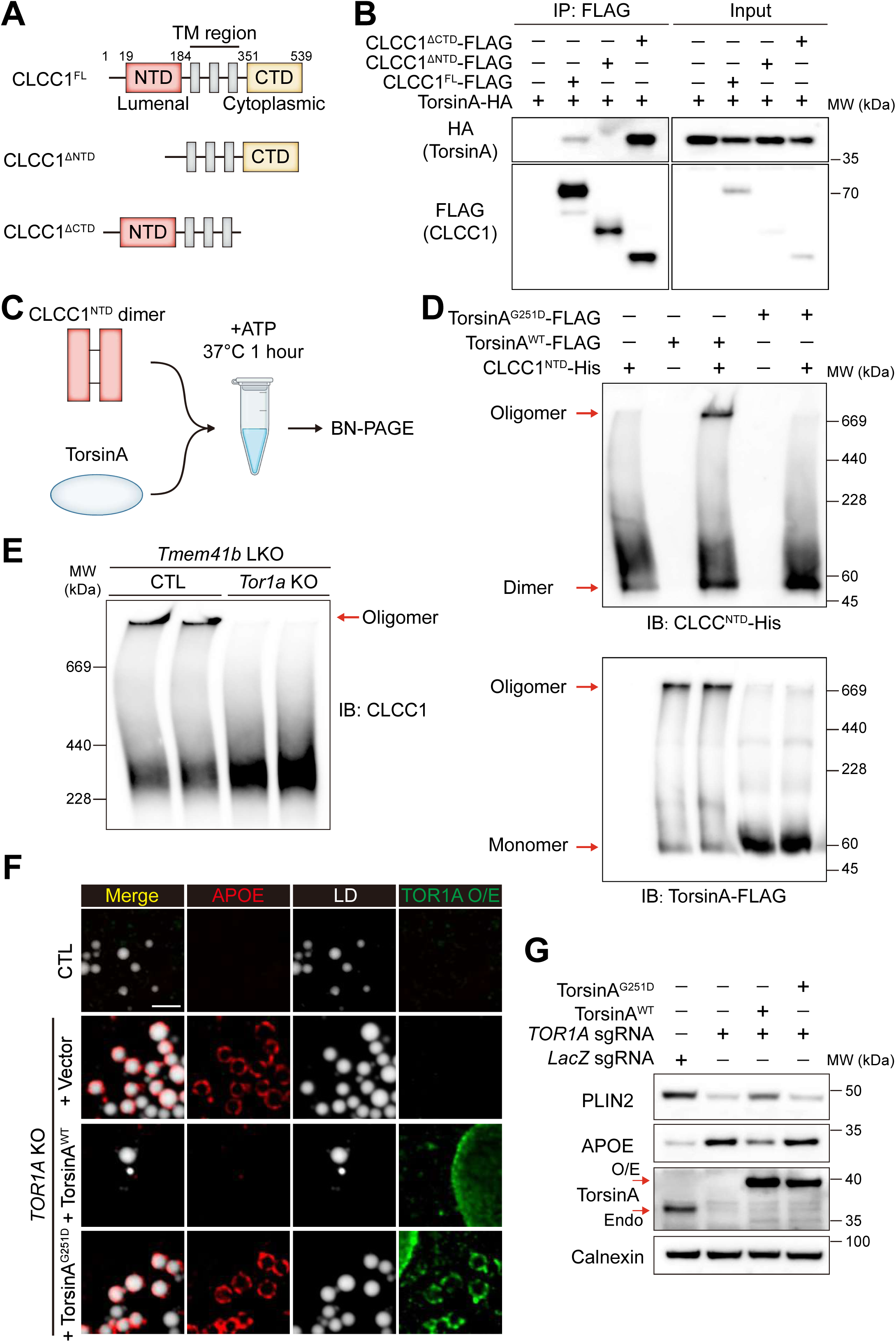
Torsins act as foldases to promote CLCC1 oligomerization in lipid control. A. Schematic of CLCC1 deletion mutants. The diagram shows the defined domains of mouse CLCC1: the N-terminal domain (amino acids 19–184), the transmembrane region (184–351), and the C-terminal domain (351–539). B. Huh7 cells were co-transduced with AAV-TBG-TorsinA and either AAV-TBG-CLCC1-FL (full-length) or the indicated CLCC1 truncation mutants for 48 hours. FLAG immunoprecipitation was performed, followed by immunoblotting with the indicated antibodies. Shown is a representative result from three independent experiments. C. Schematic of the in vitro oligomerization assay for TorsinA-mediated CLCC1 oligomerization. Reconstituted mouse CLCC1 N-terminal domain (CLCC1-NTD) dimer and human TorsinA were combined in a reaction buffer, supplemented with ATP, and incubated at 37 °C for 1 h. The reaction mixture was then analyzed by Blue Native PAGE (BN-PAGE), and oligomers were detected by immunoblotting with the indicated antibodies. D. TorsinA, but not the TorsinA^G251D^ mutant, drives the oligomeric assembly of CLCC1^NTD^ and TorsinA. CLCC1^NTD^ dimers were incubated with either TorsinA^WT^ or the TorsinA^G251D^ mutant as depicted in (C). Representative results from three independent experiments are shown. E. Loss of TorsinA ablates CLCC1 oligomerization in TMEM41B-deficient liver. BN-PAGE immunoblot analysis of CLCC1 in geLDs isolated from *Tmem41b* KO versus *Tmem41b* & *Tor1a* double KO livers (with comparable protein input) demonstrates the requirement of TorsinA for CLCC1 oligomerization. Data are representative of three experiments. F. The TorsinA^G251D^ mutant fails to rescue the geLD phenotype caused by TorsinA deficiency. *TOR1A* KO Huh7 cells were infected for 48 hours with AAV expressing luciferase (control), wild-type TorsinA-FLAG, or the G251D mutant. Cells were then fixed and co-stained with anti-APOE antibody and anti-FLAG tag antibody. Lipid droplets were visualized with LipiBlue. In contrast to TorsinA^WT^, the G251D mutant did not suppress geLD formation. Images are representative of three independent experiments. Scale bar: 1 μm. G. TorsinA^G251D^ mutant failed to rescue PLIN2 reduction and APOE accumulation induced by TorsinA deficiency. *TOR1A* KO Huh7 were infected with control AAV or AAV-TBG-TorsinA^WT^-FLAG or AAV-TBG-TorsinA^G251D^-FLAG for 48 hours prior to immunoblot with indicated antibodies. Representative results from three independent results were shown.

We thus designed an *in vitro* oligomerization assay, using recombinant Torsins and the Torsin-interacting domain of CLCC1 (Figure 4C). The inter-molecular disulfide bonds in CLCC1 N-terminal domain dimer (CLCC1^NTD^) were stabilized via expression in the reduction-defective Origami 2 *E.coli* strain (Figure S4A&B). Pulldown experiments confirmed that CLCC1^NTD^ was sufficient for TorsinA binding in an ATP-dependent manner (Figure S4C&D). BN-PAGE confirmed that Torsins oligomerize after incubation with ATP at 37°C (lane 2&3, Figure 4D, lower panel), and Torsin oligomers were enriched with CLCC1^NTD^, causing the latter to be retained at the top of the gel (lane 2&3, Figure 4D, upper panels). Importantly, a previously characterized oligomerization-defective mutant TorsinA^G251D^, failed to oligomerize and consequently failed to polymerize the CLCC1 fragment (lane 4&5, Figure 4D). In line with these *in vitro* reconstitution results, BN-PAGE of mice liver lysate showed that CLCC1 oligomerization in TMEM41B-deficient liver was abolished by further deletion of TorsinA (Figure 4E), indicating a pivotal role of Torsins in CLCC1 oligomerization.

When further examined in cells, wild type TorsinA rescued geLD induction caused by TorsinA inactivation, while the oligomerization-defective TorsinA G251D failed to rescue such defects, despite localizing to geLD-surrounding membranes (Figure 4F). Biochemical analysis further confirmed the reduction in intracellular accumulation of geLD marker ApoE by wild-type TorsinA, but not the G251D mutant (Figure 4G). These data further supported the key role of Torsin-initiated oligomerization events in the cellular function of the enigmatic ATPase.

### CLCC1 assembly rescues TorsinA deficiency in metabolic disorders

The above results suggested a critical role of Torsins in lipid regulation. Mining of public expression database revealed that TorsinA appears to be the prominent hepatic paralog in mice, while human liver also expresses Torsin3 to a moderate extent (Figure 5A), consistent with the strong lipid distortions resulted from hepatic TorsinA inactivation in mice reported previously^26^ and recapitulated in our study. Moreover, hepatic levels of TorsinA protein were significantly reduced in both genetically predisposed or diet-induced obesity models (Figure 5B, quantified in 5C), further suggesting that Torsins may become functionally compromised in metabolic diseases.

**Figure 5.**
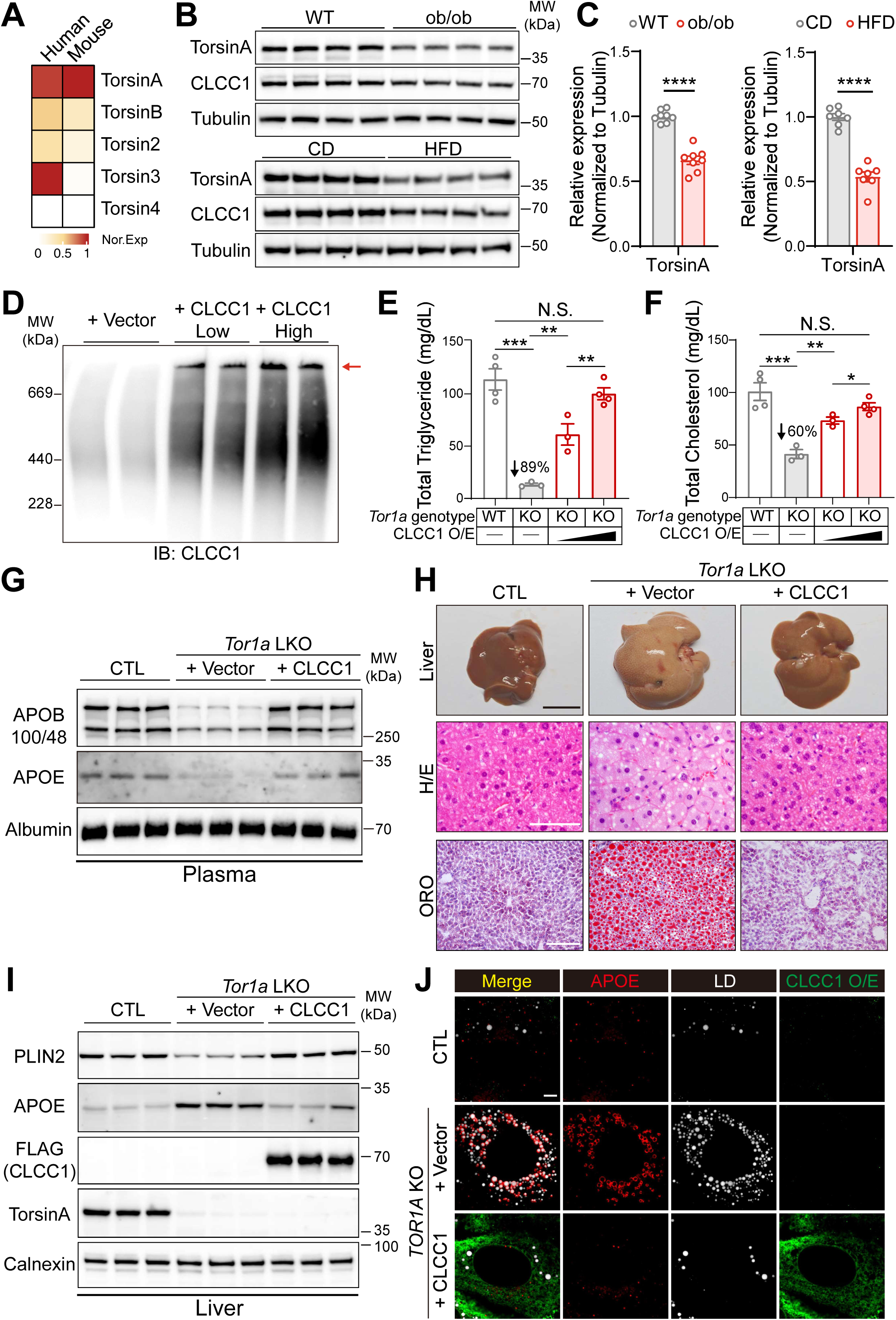
CLCC1 assembly rescues Torsin deficiency in metabolic disorders. A. Hepatic transcript levels of the Torsin family in human and mouse. Expression profiles (in Transcripts Per Million, TPM) were obtained from the GTEx (human) and EMBL-EBI Expression Atlas (mouse) databases. The normalized relative expression is shown, with the highest level within each species set to 1. B. TorsinA protein is reduced in livers from both genetic (ob/ob) and diet-induced (HFD) models of obesity. Liver lysates from the indicated mouse models were analyzed by immunoblotting with the antibodies shown. WT: age-matched wild-type mice; CD: age-matched mice on a control chow diet. C. Quantification of TorsinA protein levels from (B), showing a significant reduction in both obesity models. Group sizes: WT (n=8) vs. ob/ob (n=8); CD (n=7) vs. HFD (n=7). Data represent mean ± SEM. ****, p < 0.0001. D. Ectopically expressed CLCC1 oligomerizes in the absence of TorsinA. Hepatic TorsinA KO mice were injected with the indicated AAVs. Crude membrane fractions isolated from the livers were then analyzed by BN-PAGE and immunoblotted for CLCC1. E. CLCC1 overexpression dose-dependently rescues the depletion of plasma triglycerides (TGs) caused by TorsinA deficiency. Mice were injected with AAV-TBG-Cre alone or combined with increasing doses of AAV-TBG-CLCC1; AAV-TBG-GFP served as control. After four weeks, fasting plasma TG levels were measured. Data are mean ± SEM. N.S., not significant; **, p < 0.01; ***, p < 0.001. F. CLCC1 overexpression dose-dependently rescues the reduction in plasma total cholesterol (TC) induced by TorsinA deficiency. TC levels were measured in the plasma samples from the mice described in (E). Data are presented as mean ± SEM. N.S., not significant; *, p < 0.05; **, p < 0.01; ***, p < 0.001. G. CLCC1 overexpression rescues the reduction in plasma apolipoproteins induced by TorsinA deficiency. Plasma samples from the same mouse cohort described in (E) were analyzed by SDS-PAGE and immunoblotted with the indicated antibodies. H. CLCC1 overexpression rescues TorsinA hepatic deficiency-induced liver steatosis. Mice were sacrificed at 16 weeks post AAV injection followed by liver sectioning and indicated stains. Scale bar, 1 cm (upper panel), 100 μm (middle panel), 50 μm (lower panel). Representatives of photograph and histology were shown. I. CLCC1 overexpression rescues PLIN2 reduction and APOE accumulation induced by hepatic TorsinA deficiency. Liver lysates from mice in (H) were subjected to SDS-PAGE and immunoblotted with indicated antibodies. J. CLCC1 expression reverses geLD accumulation induced by TorsinA deficiency. Immunofluorescence of TorsinA KO Huh7 cells expressing AAV-CLCC1-FLAG or control, stained for APOE (geLDs) and LipidBlue (LDs). Data are representative of three independent experiments. Scale bar: 5 μm.

We therefore functional assess the foldase activity of Torsin in lipid control, by testing whether ectopic introduction of CLCC1 may rescue or bypass TorsinA deficiency (Figure S5A). Biochemically, ectopically expressed CLCC1 readily oligomerize in a dose-dependent manner even in the absence of TorsinA (Figure 5D). Functionally, hepatic TorsinA depleted circulating TG by ∼90%, whereas ectopic CLCC1 introduction restored such depleted plasma TG levels in a dose-dependent manner (Figure 5E). A similar rescue was also observed with plasma cholesterol levels, though the initial reduction of total cholesterol was ∼60% upon hepatic TorsinA inactivation (Figure 5F). Consistent with these lipid measurements, decreased plasma apoliproteins including ApoB and ApoE by hepatic TorsinA inactivation were also restored by ectopic introduction of CLCC1 (Figure 5G).

At the tissue level, TorsinA inactivation caused severe hepatic lipid accumulations, which were substantially rescued by CLCC1 over-expression (Figure 5H). Quantification of mice liver triglycerides and weights showed that TorsinA deficiency-induced liver steatosis were rescued by CLCC1 over-expression (Figure S5B&C). At the biochemical level, the loss of cytosolic LD marker PLIN2 and the concomitant accumulation of geLD marker ApoE caused by TorsinA inactivation were also reversed by CLCC1 introduction (Figure 5I). Lastly, cellular imaging and electron microscopy both confirmed efficient clearance of geLDs induced by TorsinA inactivation by CLCC1 expression (Figure 5J & S5D). Together, the physiological, biochemical, and cellular imaging experiment collectively demonstrated the functional bypass of TorsinA deficiency by ectopically expressed, readily-oligomerized CLCC1.

## Discussion

Since its identification as the genetic cause of early-onset dystonia, decades of work have linked Torsin ATPases to nuclear envelope integrity, ER homeostasis, and lipid metabolism^25,26,28,34^. Yet their molecular function and client substrates have remained elusive, in part because Torsins lack the canonical pore-loop features required for the unfoldase activity typical of AAA+ ATPases^32^. Here, we demonstrate that Torsins instead act as foldase-like activators of CLCC1, a recently defined regulator of ER bilayer equilibration with central roles in nuclear envelope organization and lipid flux control^17,18^. Our biochemical and cellular data reveal that Torsins promote higher-order assembly of CLCC1 oligomers. This CLCC1 oligomerization is essential for its recruitment to sites of ER bilayer imbalance, where it facilitates phospholipid equilibration^17^. Consistent with this mass-action model, ectopic CLCC1 expression bypasses the requirement for TorsinA and restores lipid secretion, ER membrane architecture, and systemic lipid homeostasis.

Our study also positions Torsins as key regulators in the trans-bilayer phospholipid equilibration at the ER, a fundamental yet poorly-understood process. Newly synthesized phospholipids arise exclusively on the cytosolic leaflet and must be efficiently redistributed to the luminal leaflet to sustain membrane biogenesis, ER remodeling, and lipoprotein assembly^35^. Moreover, constant remodeling of ER bilayer may be required for processes including ER shaping and secretory cargo export, nuclear envelope fusion, or even viral particle assembly^36–38^. The Torsin-driven CLCC1 oligomerization could therefore provide a coherent model, in which Torsins/CLCC1/TMEM41B appear to constitute a unified membrane-regulating module that ensures bilayer equilibration during high-flux lipid transport and organelle remodeling. Supporting this model, molecular dynamics simulations suggest that CLCC1 oligomers may locally destabilize the lipid bilayer (Figure 3D&E), potentially pointing to a unifying mechanism of lowering the energetic barrier for phospholipid scrambling or membrane fusion.

Although Torsins have traditionally been studied in the context of neurological disease^39^, our findings expand their roles to general cell biology and metabolic regulation. We found that these ubiquitously expressed enzymes appear to be down-regulated by metabolic stresses. The presence of multiple Torsin paralogs and enzymatic activities of these ATPases could add multiple layers of regulation, which may modulate ER bilayer homeostasis in response to specific signaling cues in distinct physiological processes^22,40,41^. Given the profound phenotypes associated with dysregulation of the fundamental control of bilayer equilibration, future therapeutic modulation of the Torsin–CLCC1–TMEM41B axis might provide new opportunities for treating dyslipidemia, MASH, and even neurological disorders. Hence, elucidating how this luminal ATPase network integrates lipid synthesis, membrane architecture, and cellular stress responses will be essential for developing strategies to restore homeostasis across diverse stress or disease settings.

## Acknowledgments

The authors thank Vivek Malhotra and Suneng Fu for critical reading of the manuscript, and acknowledge the National Center for Protein Sciences at Peking University (Beijing, China) for technical assistance with imaging. The work is supported by National Science Foundation of China (NSFC) grants 92254308, 32125021, 92157107, 92357302 and National Key R&D Program grant no. 2021YFA0804802 and 2022YFA0806502 and Key R&D Program of Zhejiang (grant no. 2024SSYS0034).

## Methods

### Mouse models

All animal housing and use procedures were approved by the Institutional Animal Care and Use Committees of Peking University, an AAALAC accredited laboratory animal facility. All mice used in the experiments were bred on the C57BL/6J background. *Tmem41b^fl/fl^* mice were generated and maintained as previously described. *Tor1a^fl/fl^* mice (Cat. NO. NM-CKO-234147) were purchased from Shanghai Model Organisms Center, Inc.. Primers for the genotypes are listed in Extended Data Table 1. C57BL/6J mice were provided by Peking University at 6 weeks of age, and ob/ob mice (RRID: IMSR_GPT: T001461) were purchased from GemPharmatech (Nanjing, China) at 4 weeks of age. Mice were housed under standardized conditions, including a temperature of approximately 22°C, a 12 h light/dark cycle, and humidity of 40%-60%. Mice had free access to food and water unless otherwise stated. Male mice aged 6-16 weeks were used in all experiments. Mice were randomly assigned to different experimental groups. For fasting studies, all experiments were performed at 9 a.m. on the second day, with fasting starting at 5 p.m. on the first day.

### Cell culture

HEK293T cells (ATCC, CRL-3216), Huh7 cells (JCRB Cell Ban, JCRB0403), and HEK293F cells (Thermo Fisher Scientific, R79007) were obtained from ATCC, JCRB, and Thermo Fisher Scientific, respectively. HEK293F suspension cells were cultured in SMM 293T-II medium (Sino Biological, M293TII), supplemented with 0.5% penicillin/streptomycin (P/S) (Caisson, PSL01) and 1% fetal bovine serum (FBS) (VisTech, SE100-011) and maintained at 37°C in a 5% CO_2_ environment, with an optimal spinner speed of approximately 100 rpm to 130 rpm. The other cell lines were cultured in Dulbecco’s modified Eagle’s medium (DMEM) (HyClone, SH30022.01B) supplemented with 1% P/S and 10% FBS; under the same temperature and CO_2_ conditions. Transfections were performed with polyethyleneimine (PEI) (Polysciences, 23966-1) according to the manufacturer’s protocol.

### DNA vector construction

AAV DNA vector (pAAV-TBG-respective cDNA) for overexpression was generated from pAAV-TBG-GFP (Addgene, 105535). In brief, respective cDNA was cloned between the NotI and BamHI restriction sites.

For fluorescence imaging, mEGFP-Perilipin2, mEGFP-CB5 (residues 90 to 124 at the carboxyl-terminal end of cytochrome B5 type A, CYB5A), were cloned between KpnI and HindIII restriction sites to substitute for enhanced GFP (EGFP) in the pAAV-CAG-EGFP vector (Addgene, 51502).

sgRNAs were designed using the Benchling platform (https://benchling.com/) to optimize editing efficiency and minimize the unintended off-target effects. The sgRNA sequences are listed in the Extended Data Table 2. Oligonucleotides were cloned into either the pX602-AAV-Cre sgRNA backbone for CRISPR/Cas9-induced acute gene knockout in mouse liver or the pLentiCRISPR V2 (Addgene, 52961) & pLentiGuide Blast for genome editing in cell lines, following established protocols as described in previous research. The pX602-AAV-Cre sgRNA construct was derived from the pX602 vector (Addgene, 61593) in our previous study. The pLentiGuide Blast construct was derived from the pLentiGuide Puro vector (Addgene, 52963), in which blasticidin S deaminase was cloned between the BsiWI and MluI restriction sites to replace puromycin N-acetyltransferase.

For protein purification, TorsinA-StrepTagII-FLAG-StrepTagII were cloned into pKH3 (Addgene, 12555) between the EcoRI and XbaI restriction sites. CLCC1-NTD were cloned into pET22b between NdeI and XhoI.

### Recombinant Adeno-Associated Virus (AAV) Production and Delivery

AAV packaging and purification were performed as previously described. For the mouse experiment, AAV shuttle plasmids, Rep/Cap (2/8) plasmids, and helper plasmids were transfected into HEK293T cells using PEI. After 60 h post transfection, cells were harvested and virus was purified, quantified by both Coomassie Blue R250 staining and qPCR.

AAV-2/8 was administered by tail vein injection. To achieve acute inactivation of the hepatic *Tor1a* gene, each *Tor1a^fl/fl^* mouse was randomly administered with AAV-TBG-GFP or AAV-TBG-CRE at a viral genome copy number of 1×10^11^. For rescue experiments, each *Tor1a^fl/fl^* mouse was simultaneously injected with 1×10^11^ viral genome copies of AAV-TBG-CRE and 1×10^10^ or 4×10^10^ viral genome copies of AAV-TBG-mCLCC1-StrepTagII-FLAG-StrepTagII.

For AAV delivery into a cultured cell line, pRep/Cap (2/8) was replaced by pAAV-DJ for AAV-DJ preparation. AAV-DJ with 1×10^10^ genome copies number was used per well in 6-well plate to delivery indicated gene into cultured cells. Subsequent experiments were conducted 24 h after AAJ-DJ infection, unless otherwise specified.

### Lentiviral packaging and knockout/knockin cell construction

HEK293T cells were used for lentivirus packaging. Briefly, the lentivirus shuttle plasmids, psPAX2 (Addgene, 12260), and pMD2.G (Addgene, 12259) plasmids were introduced into the cells using PEI according to the protocol provided by the manufacturer. Fouty-eight hours post transfection, the medium containing the lentivirus was collected and subsequently added to the WT Huh7 cells. Transducted cells were selected by antibiotic at 48 h after infection.

For control, *TMEM41B* KO, *CLCC1* KO and *TOR1A* KO Huh7 cell construction, pLentiCRISPR V2 lentivirus containing *LacZ* targeting sgRNA, human *TMEM41B* targeting sgRNA, human *CLCC1* targeting sgRNA and human *TOR1A* targeting sgRNA (supplementary table 1) were generated and infected into Huh7 cells, respectively. Two days after transduction, cells were selected with puromycin (Sigma, P8833). For TMEM41B & TorsinA double deficient Huh7 cells, *TMEM41B* KO Huh7 cells were infected with pLentiGuide Blast lentivirus containing control or human *TOR1A* targeting sgRNA, and further selected with blasticidin (Wako, 022-18713).

For TOR1A knock-in Huh7 cells construction, a twin StrepTagII and HaloTag were inserted into the C-terminus of human TorsinA via CRISPR/Cas9 system. In brief, sgRNA (Supplementary table) along with homology-directed repair (HDR) donor template were delivered into the Cas9 stably expressed Huh7 cells via an AAV vector. The HDR donor template, flanked by AAV inverted terminal repeats (ITRs), includes one 1000 bp homology arms upstream and one 1000 bp homology arms downstream of the DSB site.

### Immunoblotting

Liver lysates were prepared using RIPA buffer supplemented with protease inhibitors. Protein concentrations were measured using the BCA Protein Assay Kit. 1× SDS/PAGE sample buffer was added. Equal amounts of protein were loaded onto 3% to 15% Tris-acetate SDS/PAGE gels and transferred overnight at 4°C to nitrocellulose membranes (Cytiva, 10600006). The membranes were then blocked with 5% milk in TBST□(20 mM Tris pH 7.4, 150 mM NaCl and 0.1% Tween 20) for 1 h at room temperature. Primary antibodies were diluted in TBST containing 5% milk and 0.02% Na_3_N, incubated with membranes overnight at 4°C and washed 3 times with TBST at room temperature, for 5 min each time. HRP-conjugated goat secondary antibodies were diluted in TBST containing 5% milk and incubated for 1 h at room temperature, followed by 3 washes. Membranes were visualized after exposure to ECL substrate (Thermo Fisher, 34578).

The following primary antibodies were used for immunoblotting: rabbit polyclonal anti-mTorsinA(abcam, ab34540,1:1000); rabbit polyclonal anti-hTorsinA (Proteintech,10296-1-AP,1:1000); rabbit polyclonal anti-TorsinB (Proteintech, 21945-1-AP, 1:500); rabbit monoclonal anti-FLAG (Proteintech, 66008-4-Ig, 1:5000); mouse monoclonal anti-His (Proteintech, 66005-1-Ig, 1:5000); rabbit polyclonal anti-Perilipin2 (Proteintech, 15294-1-AP, 1:1,000); rabbit polyclonal anti-Perilipin2 (Cell Signaling Technology, 45535, 1:1,000); rabbit monoclonal anti-hApoE (abcam, ab52607, 1:1000); rabbit polyclonal anti-Calnexin (Proteintech, 10427-2-AP, 1:1,000); rabbit polyclonal anti-ApoB (Proteintech, 20578-1-AP, 1:1,000); rabbit polyclonal anti-mApoE, (Fitzgerald, 10R-10633, 1:1,000); rabbit polyclonal anti-Tubulin (Proteintech, 11224-1-AP, 1:1,000); homemade rabbit polyclonal antibody against the C-terminal epitope of mouse CLCC1 (residues 351-539, 1:5,000).

The following secondary antibodies were used: goat anti-mouse IgG (H+L) secondary antibody (Thermo Fisher, 31430, 1:10,000) and goat anti-rabbit IgG (H+L) secondary antibody (Thermo Fisher, 31460, 1:10,000).

### Immunoprecipitation (IP) and quantitative mass-spectrometry

For tandem IP purification of mCLCC1-StrepTagII-FLAG-StrepTagII (SFS) in mice liver, 2E9/4E9 viral genome copies (GC) of AAV-TBG-mCLCC1-SFS were delivered to *Tmem41b^fl/fl^* mice together with AAV-TBG-GFP/Cre for WT/KO group, respectively. At three weeks post AAV injection, mice were sacrificed and harvested the liver. For each purification, 1 g of liver tissue was lysed in cell lysis buffer (50 mM Tris-HCl, pH 7.4, 150 mM NaCl, 1% NP-40, 10% glycerol with protease inhibitor, CLB) using a Dounce homogenizer. The lysate was sequentially centrifuged at 1,000g for 10 min, 13,000g for 10 min and 150,000g for 30min. The supernatant was incubated with ProteinA beads to remove non-specifically binding proteins. Then the supernatant was mixed with anti-FLAG beads for 2 hours at 4 □ to enrich CLCC1, followed by 4 washes with CLB and elution with 3×FLAG peptide. The eluate was further incubated with Streptactin beads for 2 hours at 4 □. Finally, the Streptactin beads were washed 10 times with CLB and eluted with 5 mM dethiobiotin in CLB. Eluted proteins were separated by SDS-PAGE and visualized by silver staining using a commercial kit.

Quantitative mass-spectrometry was used to identify the CLCC1 complex in liver tissue from WT and TMEM41B KO mice. Proteomic analysis was performed as previously described (ref). Proteins identified with at least one unique peptide were retained for further analysis. Significantly altered proteins were defined as those showing a fold change > 10 and a *p*-value < 0.05.

### Immunofluorescence and confocal microscopy

Cells were seeded on 22-mm glass coverslips (Thermo Fisher, 12-541-007) and grown to 40-60% confluence prior to processing. After 24–48 hours, cells were washed with PBS and fixed with 4% paraformaldehyde (PFA) in PBS for 15 minutes at room temperature (RT). Following fixation, cells were washed three times with PBS and permeabilized with Immunostaining Permeabilization Buffer containing Saponin (Beyotime, P0095) for 15 minutes at RT. After permeabilization, cells were washed three times with PBS and blocked with 2.5% goat serum in PBS for 1 hour at RT. Subsequently, cells were incubated with primary antibodies diluted in blocking buffer overnight at 4°C. The next day, cells were washed three times with PBS (15 minutes per wash) at RT, followed by incubation with fluorescence-conjugated secondary antibodies for 1 hour at RT. After secondary antibody incubation, cells were washed three times with PBS (20 minutes per wash) at RT. For lipid droplet staining, cells were incubated with Lipi-Blue (DOJINDO, LD01; diluted 1:100 in PBS) for 30 minutes and then washed with PBS. Finally, coverslips were mounted on glass slides using approximately 15 µL of Fluorescence Mounting Medium (DAKO, S196430), ensuring the cell side faced the mounting medium.

The following primary antibodies were used for immunofluorescence: mouse monoclonal anti-FLAG (M2) (Sigma, F1804, 1:1000); homemade rabbit polyclonal antibody against the C-terminal epitope of mouse CLCC1 (residues 351-539, 1:500); rabbit monoclonal anti-ApoE (abcam, ab52607, 1:100); mouse monoclonal anti-ApoE (Cell Signaling Technology, 74417, 1:100)

The following secondary antibodies were used for immunofluorescence: Goat anti-Rabbit IgG (H+L) Cross-Adsorbed Secondary Antibody, Alexa Fluor^TM^ 488 (Thermo Fisher, A-11008, 1:1,000); Goat anti-Rabbit IgG (H+L) Cross-Adsorbed Secondary Antibody, Alexa Fluor^TM^ 568 (Thermo Fisher, A-11011, 1:1,000); Goat anti-Mouse IgG (H+L) Cross-Adsorbed Secondary Antibody, Alexa Fluor^TM^ 488 (Thermo Fisher, A-11001, 1:1,000); Goat anti-Mouse IgG (H+L) Cross-Adsorbed Secondary Antibody, Alexa Fluor^TM^ 568 (Thermo Fisher, A-11004, 1:1,000)

Images were acquired on a spinning disk confocal SpinSR10 (EVIDENT) and processed with ImageJ (NIH).

### HaloTag staining and imaging

Torsin1A^Strep-Halo^ knock-in cells were seeded on 22-mm glass coverslips (Thermo Fisher, 12-541-007). For labeling, cells were incubated with 10 nM Janelia Fluor® JFX_554_ HaloTag® Ligand (Promega, HT103A) diluted in culture medium for 30 minutes at 37□°C under 5% CO□ in a cell culture incubator. After labeling, cells were washed three times with fresh culture medium under the same incubator conditions. Finally, cells were fixed with 4% PFA for subsequent immunofluorescence staining.

### EM samples preparation and images

Adult mice were transcardially perfused with 100 mM sodium phosphate buffer (PB, pH 7.4), followed by pre□fixation using a solution containing 2.5% glutaraldehyde and 0.8% paraformaldehyde. The liver tissues were then dissected and immersed in the same pre□fixation solution for 2□h at room temperature, before being trimmed into smaller pieces. Samples were further fixed overnight at 4□°C in the pre□fixation solution. Following rinses in PB, the tissues were treated with 0.1□M imidazole in 0.1□M PB for 30□min and post□fixed in 2% osmium tetroxide prepared in 0.1□M PB. After thorough washing with ultrapure water, the tissues were stained with 1% uranyl acetate at 4□°C overnight. Dehydration was performed using a graded acetone series, followed by embedding in epoxy resin and polymerization at 60□°C for 24□h. Ultrathin sections of approximately 60□nm were cut with a Leica EM UC7 ultramicrotome, collected on copper grids, and examined using an FEI Tecnai G2 20 Twin transmission electron microscope equipped with a CMOS camera (EMSIS, Germany). Evaluation of liver tissue structure was conducted under double□blind conditions.

### Isolation of subcellular organelles

Mice were fasted for 4 h. After anesthesia, the liver was perfused through the portal vein to remove blood, excised, and cut into approximately 1 mm^3^ pieces in Buffer A (25 mM Tricine pH 7.6, 250 mM sucrose) plus 0.2 mM PMSF and protease inhibitors (Roche, 4693132001). Tissue pieces were homogenized 20 times on ice using a Dounce type glass-Teflon homogenizer. The lysate was centrifuged twice at 500g for 10□min at 4□°C to remove debris and nuclei, the resulting post□nuclear supernatant (PNS) was collected. The PNS was then centrifuged at 8,000 g for 30 min twice to remove mitochondria. The supernatant was further centrifuged at 100,000 g for 30 min. After this step, the pellet was desiganeted as the ER fraction for further analysis and the white band of LD floating on the top of the solution was collected as the geLD fraction.

For IB analysis, both ER and geLD fractions were lysed in RIPA buffer to extract proteins. A total of 10 µg protein per sample was separated by SDS-PAGE and analyzed by IB. For BN-PAGE, experiment procedures are described below.

### BN-PAGE

Blue native-PAGE (BN-PAGE) analysis of CLCC1 assembly was performed as previously described. In brief, the geLD fraction or membrane fraction from mice of the indicated genotypes was incubated with Buffer D (50 mM Imidazole/HCl pH 7.0, 50 mM NaCl, 2 mM 6-Aminohexanoic acid, 10% glycerol, 1% n-dodecyl-ß-D-maltoside (DDM, Qisong biological, QS81007015) and protease inhibitors) for 1h at 4□°C, followed by centrifugation at 100,000g for 30 min. Protein concentrations in the supernatants were measured using the BCA Protein Assay Kit. Coomassie Brilliant Blue G-250 dye (5% stock) was added at a final dye-to-detergent ratio of 1:8 (g/g). Equal amounts of protein were loaded onto a 3-10% blue native-PAGE gel, transferred to a polyvinylidene difluoride (PVDF) membrane (Cytiva, 10600023) at 20 V for 3 h, and subsequently analyzed by IB.

### Protein expression and purification

For TorsinA-WT and the TorsinA-G251D variant, HEK293F cells were transfected with pKH3-TorsinA-StrepTagII-FLAG-StrepTagII plasmids. After 48 h, cells were harvested by centrifugation at 1,000g for 10 min at 4 °C, then washed with ice-cold TBS buffer. Cell pellets were resuspended in Buffer A (50□mM Tris-HCl pH□7.4, 150□mM NaCl, 5 mM MgCl_2_, 1% DDM, 10% glycerol, supplemented with protease inhibitors). After 2 h incubation, the cell lysate was centrifuged at 100,000g for 30 min, and the supernatant was incubated with pre-washed anti-DYKDDDDK affinity beads (Smart-lifesciences, SA042500) for 2 h at 4 °C, then beads were washed with Buffer B (50□mM Tris-HCl pH□7.4, 150□mM NaCl, 5mM MgCl_2_, 0.05% DDM and 10% glycerol). The proteins were eluted with Buffer A containing 0.4 mg/mL 3XFLAG peptide. The peptide-eluted products were further incubated with Streptactin beads 4FF (Smart-lifesciences, SA053500) for 2 h at 4 °C. After washing, proteins were eluted with Buffer A containing 5 mM desthiobiotin (Sigma, 71610-M). Finally, the eluted proteins were concentrated and buffer-exchanged into

Buffer B using Amicon Ultra 0.5 mL centrifugal filters (10 kDa cutoff, Millipore, UFC501008) and quantified by Coomassie Blue R250 staining using BSA standards.

For CLCC1-NTD-6×His dimer purification, Origami2 (DE3) pLysS cells were used for expression by pET22B vector. The cells were grown at 37 □ in DYT medium (1.6% (wt/vol tryptone, 1% yeast extract, and 0.5% NaCl, pH 7.0) supplemented with 100 μg/mL ampicillin to OD_600_ of 0.8 and protein expression was induced with 1 mM isopropyl-β-D-thiogalactopyranoside (IPTG) at 18 □ overnight. The cells were collected by centrifugation at 6,000×g for 10 min and the pellets were resuspended in PBS and centrifuged at 6,000×g for 10 min to remove residual medium. Then the pellets were snap-frozen in liquid nitrogen and stored at -80 □. For protein purification, the cells were lysed by sonication in lysis buffer (20 mM Tris-HCl, pH 7.5, 500 mM NaCl and 10 mM imidazole, supplemented with protease inhibitors). After centrifugation of 14,000×g for 30 min at 4 □, the supernatant was incubated with 1 mL Ni Sepharose resin at 4 □ for 2 hours. Unbounded materials were washed away with 20 mL of lysis buffer. The CLCC1-NTD-6×His protein was then eluted using 5 mL of elution buffer (20 mM Tris-HCl, pH 7.5, 500 mM NaCl and 500 mM imidazole). The eluate was concentrated to ∼1 mL using an ultrafilter tube and loaded onto a Resource Q column. The proteins were eluted with a linear NaCl gradient from 50 mM to 1 M. Fractions containing the CLCC1-NTD dimer were pooled and further purified by size-exclusion chromatography on a Superdex 75 HiLoad column. Finally, for long-term storage, glycerol was added to the protein solution to a final concentration of 10% (vol/vol), followed by snap-freezing in liquid nitrogen and storage at –80 °C.

### *In vitro* oligomerization assay

Two μg of purified CLCC1-NTD were incubated in 120 μL oligomerization buffer (50 mM Tris-HCl, pH 7.4, 30mM NaCl, 50 mM NaAc, 5 mM MgCl_2_, 10% glycerol, 2mM ATP, 0.05% DDM) at 37 °C for 1 h, with or without 1 ug TorsinA-WT or TorsinA-G251D. Then the reaction mixture was subjected to centrifugation at 13,000 g for 10 min. The resulting supernatant was split equally, with one portion applied to BN-PAGE analysis and the other to pulldown assays.

### Plasma characterization

Blood samples were collected from the tail tips of mice fasted for 16 h using a heparinized capillary. Plasma was separated by centrifugation at 6000 rpm for 10 min at 4°C. Triglyceride and total cholesterol levels were determined using specific commercial kits (Sigma, TR0100; 000180 from Zhongsheng beikong, respectively) according to the manufacturer’s instructions.

### Histology

Tissue samples were collected and preserved in 4% paraformaldehyde (PFA) in PBS (Leagene, DF0135). Tissue embedding, sectioning, and hematoxylin and eosin (H&E) staining were performed by the Pathology Center of Peking University. For oil red O staining, tissues were embedded in OCT compound and rapidly frozen. Cryosections of 8 μm thickness were cut and stained with Oil Red O according to the manufacturer’s instructions.

### Quantification of hepatic triglycerides

Liver samples were quantified and homogenized in PBS. Lipids were extracted from the homogenates according to the established protocols of the modified Bligh-Dyer method. Briefly, the homogenates were vigorously mixed with a chloroform-methanol mixture (2:1). After centrifugation, the organic phase was carefully collected and concentrated using a rotary evaporator under vacuum. The lipid extract obtained was reconstituted in a solution of 15% Triton X-100 (Sigma, X100) in ddH_2_O. Quantification of triglycerides was performed as previously described.

### Alphafold prediction

Based on predictions that CLCC1 forms ring-like oligomers^17,18,38^, with the highest confidence (ipTM) for a hexadecamer, we used AlphaFold3^42^ to model its structure with 16 copies. Concurrently, given evidence that Torsins may adopt hexameric assemblies^20^, we predicted a complex using these 16 copies of mouse CLCC1 (residues 19-351, lacking the C-terminal domain) alongside 6 copies of full-length mouse TorsinA. The overall ipTM score of the shown TorsinA/CLCC1 oligomer is 0.36.

### Coarse grained molecular dynamics simulations

Coarse grained molecular dynamics simulations of the AlphaFold3 predicted model of CLCC1-Torsin complex were prepared with the CHARMM-GUI Martini Bilayer Builder with the Martini 2.2 forcefield with elastic networks imposed (elnedyn22) and carried out using GROMACS 2023^43,44^. The model was embedded in a 25 × 25 nm DOPC bilayer, and the system was solvated, neutralized and ionized using 150 mM NaCl. Following the standard CHARMM-GUI protocol, the system was subjected to a short energy minimization, followed by a 5-step equilibration protocol spanning 4.75 ns in total in which restraints on the protein (1000 kJ mol^-1^ nm^-2^, 500 kJ mol^-1^ nm^-2^, 250 kJ mol^-1^ nm^-2^, 100 kJ mol^-1^ nm^-2^, 50 kJ mol^-1^ nm^-2^) and lipid headgroups (200 kJ mol^-1^ nm^-2^, 100 kJ mol^-1^ nm^-2^, 50 kJ mol^-1^ nm^-2^, 20 kJ mol^-1^ nm^-2^, 10 kJ mol^-1^ nm^-2^) were gradually reduced. Production simulations were carried out in the NPT ensemble, with a pressure of 1 bar maintained using a Parinello-Rahman barostat with semi-isotropic conditions^45^. The temperature was maintained at 310 K using a v-rescale thermostat^46^. Electrostatics were treated using the reaction-field method with a dielectric constant of 15, and a Coulomb cutoff of 1.1□nm. A cutoff of 1.1 nm was applied for van der Waals interactions. To prevent unrealistic protein deformation during the simulation, we imposed weak backbone position restraints of 5 kJ mol^-1^ nm^-2^ in production simulations. Production simulations were carried out in triplicate for 5 μs using a timestep of 20 fs.

Simulations were visualised and figures produced using VMD^47^. DOPC lipid order parameters were calculated across a concatenated trajectory containing all three replicates aligned to the protein backbone in which each frame represents 10 ns of simulation using the gorder tool^48^.

### Quantification and statistical analysis

Unless otherwise specified in the figure legends, all data are expressed as mean ± SEM. No statistical method was used to predetermine sample size. Statistical analyses were conducted using GraphPad Prism 9, with Student’s t-test as detailed in the figure legends. A P value of less than 0.05 was considered statistically significant, with significant levels marked accordingly in the figures. Results shown are representative of at least three independent experiments. For mouse studies, animals were randomly assigned to groups. Imaging and histological assessments were performed in a blinded manner.

---

**Supplementary Figure 1.**
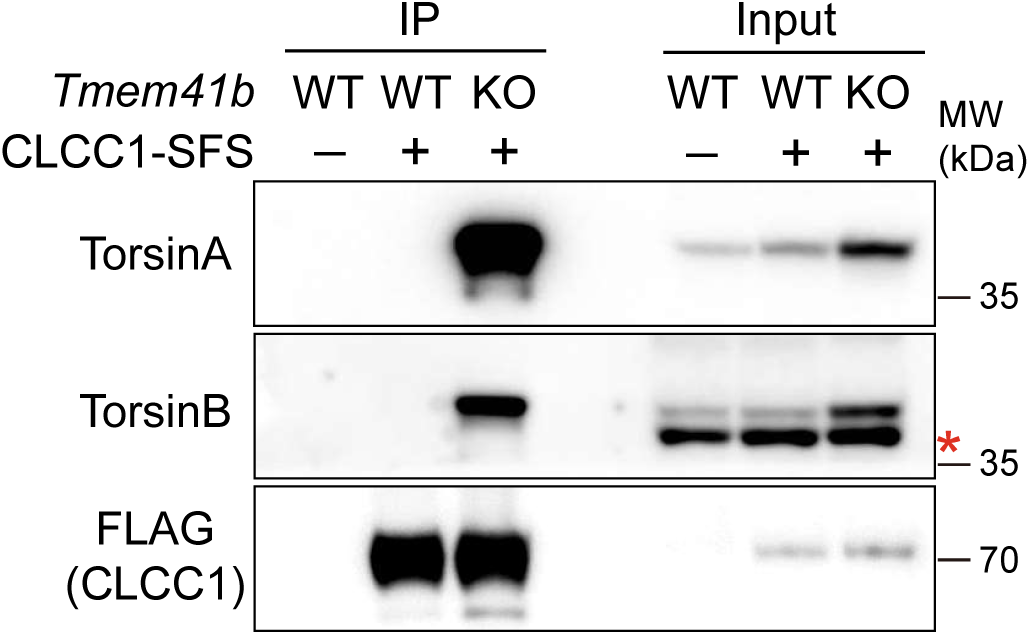
Related to Figure 1. identification of Torsins as inducible partners of CLCC1. The inducible interaction between CLCC1 and Torsins was confirmed by co-immunoprecipitation (co-IP). CLCC1 complexes were purified from livers of the indicated genotypes, subjected to SDS-PAGE, and immunoblotted with the indicated antibodies. The asterisk indicates a non-specific band.

**Supplementary Figure 2.**
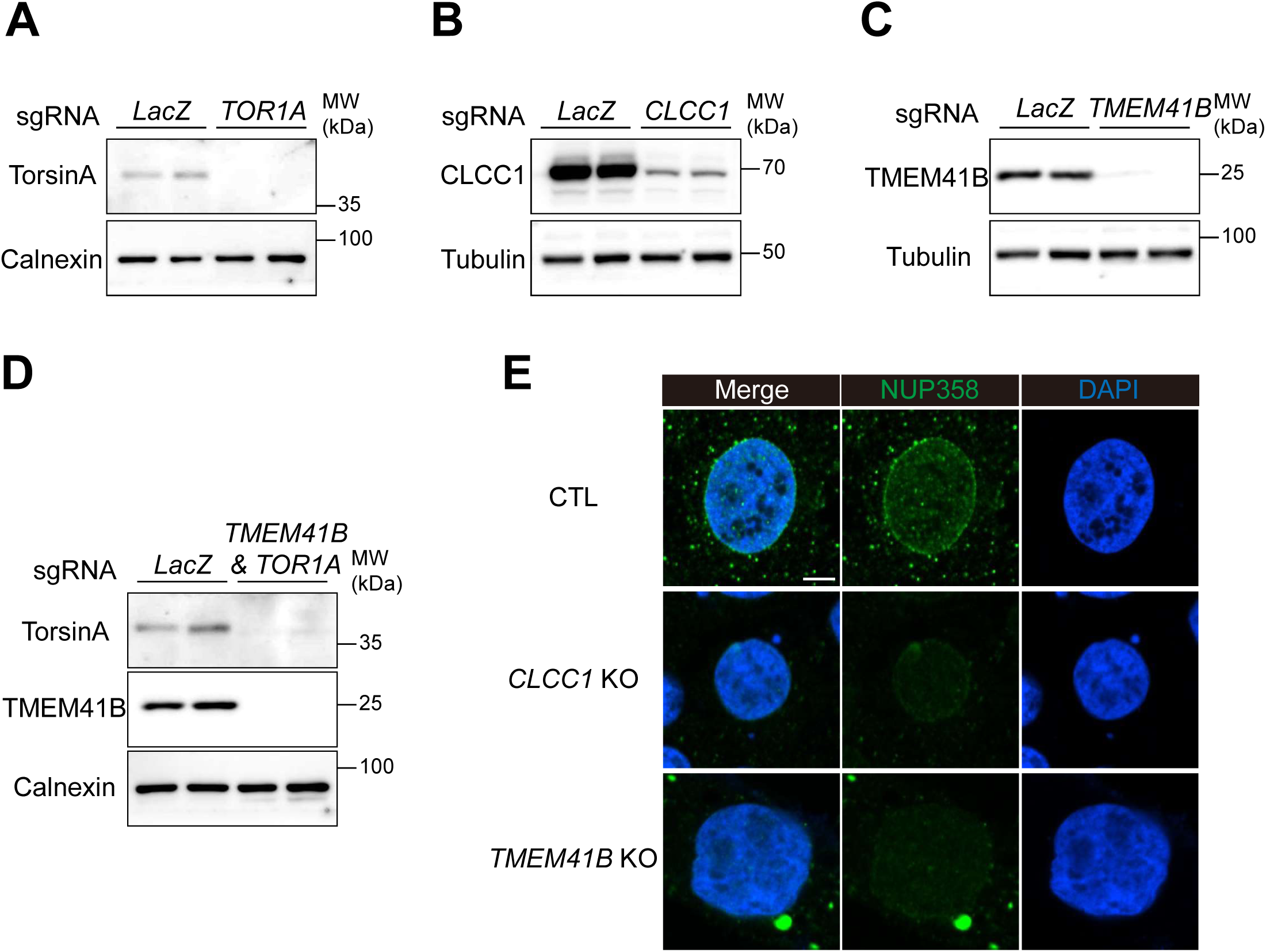
Related to Figure 2. Torsin deficiency induces lipid mis-deposition across the ER membrane. (A) Verification of *TOR1A* knockout. Lysates of *LacZ* or *TOR1A* sgRNA expressed Huh7 cells were subjected to SDS-PAGE and immunoblotted with the indicated antibodies. (B) Verification of *CLCC1* knockout. Experiment was done as in (A). (C) Verification of *TMEM41B* knockout. Experiment was done as in (A). (D) Verification of *TOR1A & TMEM41B* double KO. Experiment was done as in (A). (E) Disrupted cellular NUP358 localization induced by *CLCC1* or *TMEM41B* KO in Huh7 cells. CTL, *CLCC1* KO and *TMEM41B* KO Huh7 cells were fixed and immuno-stained with anti-NUP358 antibody. DAPI was used for labeling the nucleus. Scale bar, 5 μm.

**Supplementary Figure 3.**
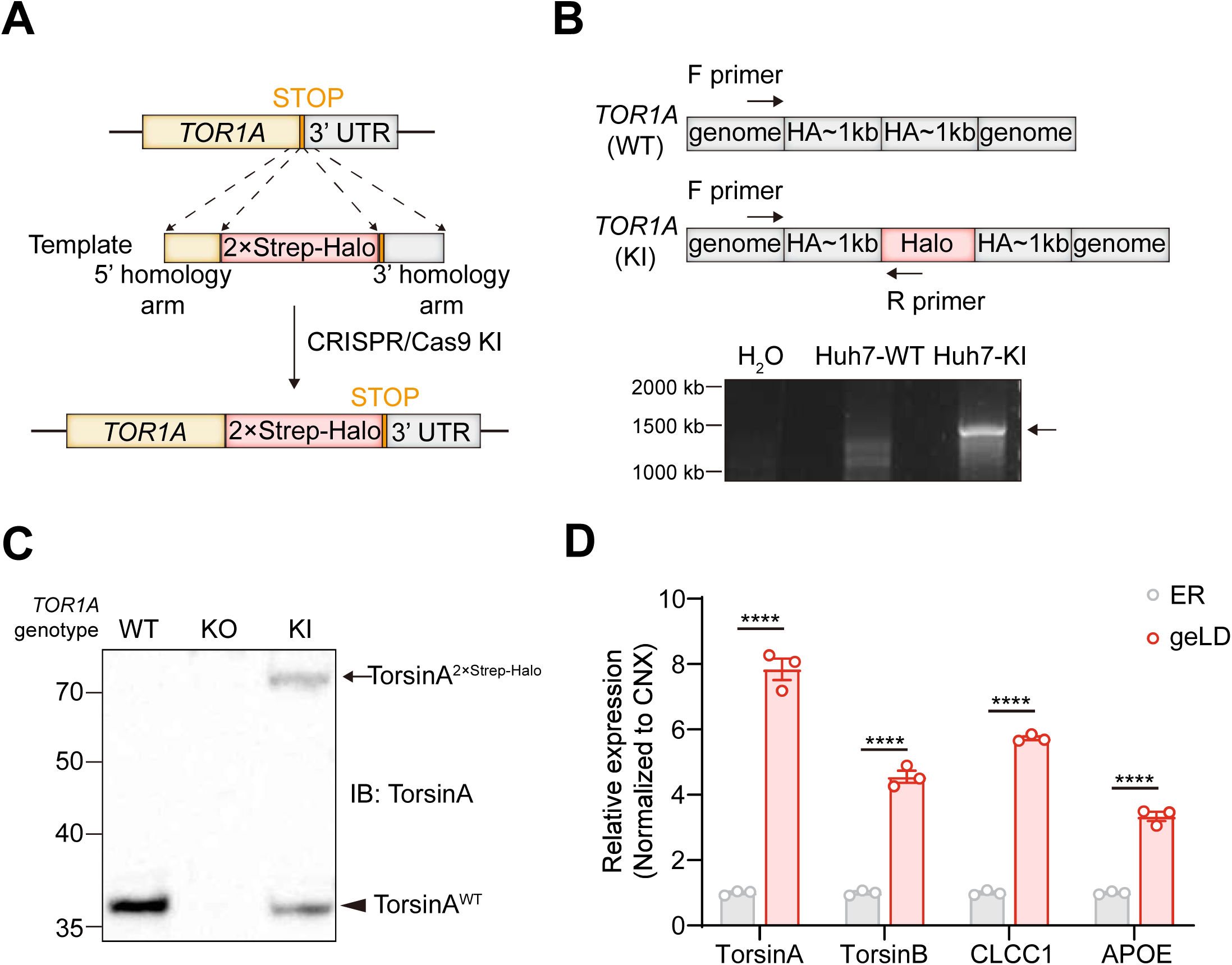
Related to Figure 2. Torsin deficiency induces lipid mis-deposition across the ER membrane. (A) Schematic for generating TorsinA knock-in (KI) Huh7 cells. A twin-Strep-Halo tag was inserted at the C-terminus of the *TOR1A* gene via CRISPR/Cas9-mediated genome editing. (B) Schematic and genotyping results of the *TOR1A*-2×Strep-Halo knockin. Genotyping PCR was performed using a forward primer upstream of the 5’ homology arm and a reverse primer within the inserted Halo tag sequence. (C) Validation of *TOR1A*-2×Strep-Halo knockin at the protein level. Lysates from control (CTL), *TOR1A* knockout (KO), and *TOR1A* knockin (KI) Huh7 cells were analyzed by SDS-PAGE and immunoblotting with an anti-TorsinA antibody. (D) Quantification of the immunoblot in Figure 2E. ****, p < 0.0001.

**Supplementary Figure 4.**
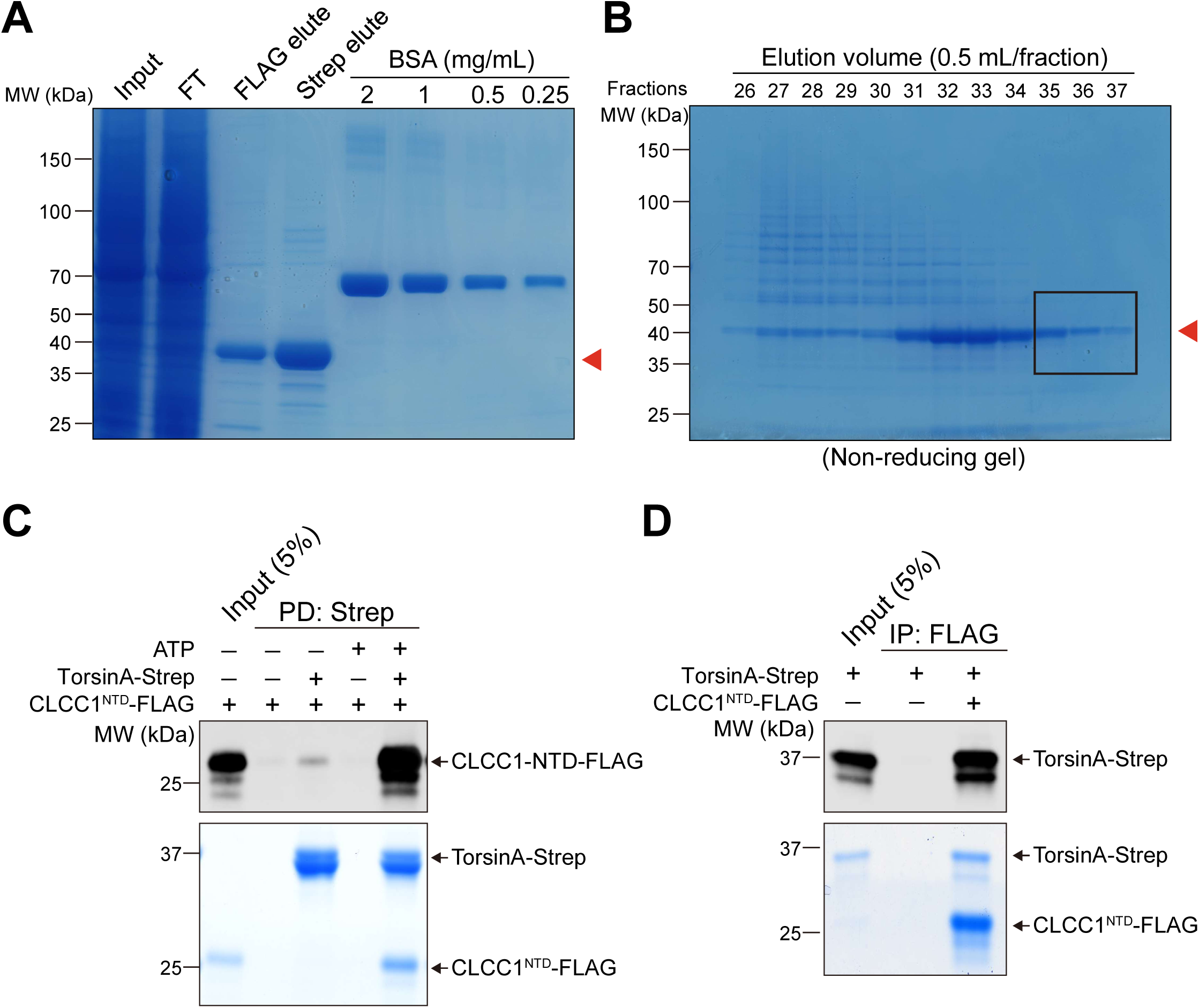
Related to Figure 4. Torsins act as protein foldases to promote CLCC1 oligomerization in lipid control. (A) Purification of human TorsinA-Strep-FLAG-Strep. TorsinA-Strep-FLAG-Strep was expressed in 293F cells and purified via tandem immunoprecipitation. Aliquots of the indicated fractions were analyzed by SDS-PAGE and Coomassie blue staining. The red arrowhead indicates the purified TorsinA-Strep-FLAG-Strep protein. (B) Purification of the mouse CLCC1-NTD dimer. CLCC1-NTD-6×His was expressed in Origami2 cells and purified using Ni Sepharose resin, followed by sequential FPLC on Q and Superdex 75 columns. Equal volumes of peak fractions were analyzed by non-reducing SDS-PAGE with Coomassie blue staining. The red arrowhead indicates the CLCC1-NTD dimer. (C) ATP-dependent interaction between purified TorsinA-Strep and CLCC1^NTD^-FLAG. Reconstituted CLCC1^NTD^-FLAG was incubated with or without purified TorsinA, in the presence of ATP, followed by Strep pulldown. Representative result from three independent experiments was shown. (D) Reciprocal interaction between purified TorsinA-Strep and CLCC1^NTD^-FLAG. Reconstituted TorsinA was incubated with or without purified CLCC1^NTD^-FLAG, in the presence of ATP, followed by FLAG IP. Representative result from three independent experiments was shown.

**Supplementary Figure 5.**
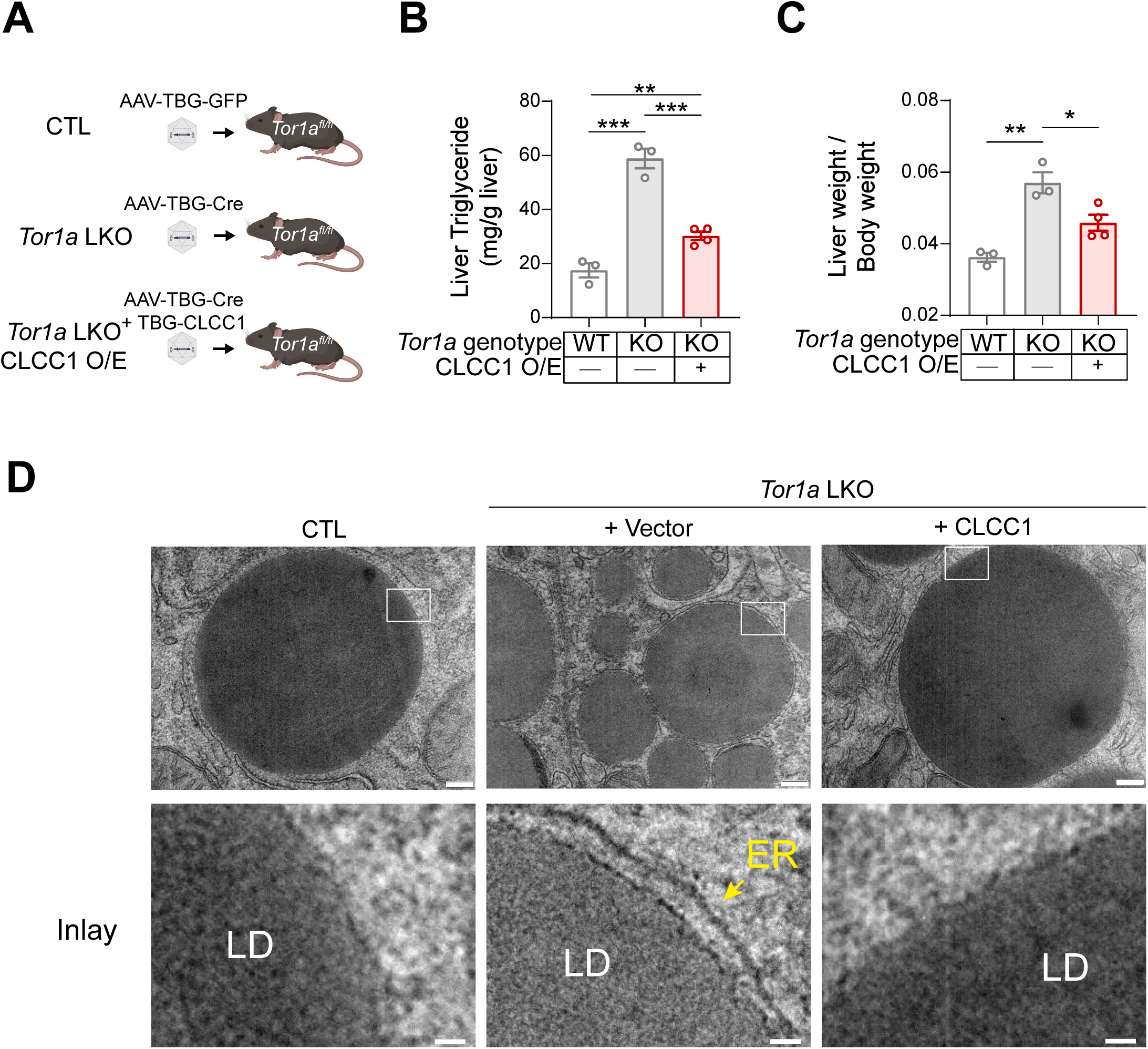
Related to Figure 5. CLCC1 assembly rescues TorsinA deficiency in metabolic disorders. (A) Schematic of the rescue strategy by CLCC1 overexpression in TorsinA KO mice. To counteract the phenotype induced by hepatic TorsinA knockout, 6-week-old *Tor1a* flox mice were injected with AAV-TBG-Cre (for knockout) together with either AAV-TBG-CLCC1-FLAG (rescue) or AAV-TBG-GFP (control). (B) Quantification of liver triglycerides (TGs) in the mice from (A). Mice were euthanized at 16 weeks post-AAV injection for analysis. Livers were dissected, and lipids were extracted from homogenates for TG quantification. **, p < 0.01; ***, p < 0.001. (C) Quantification of liver weight. Mice from the experimental groups in (A) were euthanized at 16 weeks post-AAV injection, and livers were collected and weighed. *, p < 0.05; **, p < 0.01. (D) CLCC1 overexpression rescues TorsinA deficiency-induced geLDs. Mice from (A) were transcardially perfused with PBS followed by fixative and processed for transmission electron microscopy (TEM) imaging. The endoplasmic reticulum (ER) is indicated by yellow arrowheads. Scale bars: 200 nm (upper), 30 nm (lower).

